# Molecular and morphological circuitry of the octopus sucker ganglion

**DOI:** 10.1101/2025.02.10.637560

**Authors:** Cassady S. Olson, Aashna Moorjani, Clifton W. Ragsdale

**Affiliations:** Committee on Computational Neuroscience, University of Chicago, Chicago, IL 60637; Department of Neurobiology, University of Chicago, Chicago, IL 60637

**Keywords:** sensorimotor control, neural circuitry, *Octopus bimaculoides*, cephalopod, invertebrate, muscular hydrostat, neuroethology

## Abstract

The octopus sucker is a profoundly complex sensorimotor structure. Each of the hundreds of suckers that line the octopus arm can move independently or in concert with one another. These suckers also contain an intricate sensory epithelium, enriched with chemotactile receptors. Much of the massive nervous system embedded in the octopus arm mediates control of the suckers. Each arm houses a large axial nerve cord (ANC), which features local enlargements corresponding to each sucker. There is also a sucker ganglion, a peripheral nervous element, situated in the stalk of every sucker. The structure and function of the sucker ganglion remains obscure. We examined the cellular organization and molecular composition of the sucker ganglion in *Octopus bimaculoides*. The sucker ganglion has an ellipsoid shape and features an unusual organization: the neuropil of the ganglion is distributed as a cap aborally (away from the sucker) and a small pocket orally (towards the sucker), with neuronal cell bodies concentrated in the space between. Using in situ hybridization, we detected positive expression of sensory (*PIEZO*) and motor (*LHX3* and *MNX*) neuron markers in the sucker ganglion cell bodies. Nerve fibers spread out from the sucker ganglion, targeting the surrounding sucker musculature and the oral roots extending to the ANC. Our results indicate that the sucker ganglion is composed of both sensory and motor elements and suggest that this ganglion is not a simple relay for the ANC but facilitates local reflexes for each sucker.

## Introduction

The octopus arm is a remarkable appendage, famously able to contort in near infinite degrees of freedom (Hochner et al., 2023; Olson and Ragsdale, 2023). The hundreds of suckers that line each arm are equally remarkable. These suckers are vital for the survival of the octopus, allowing them to catch prey, distinguish food from other objects, pass food to and from the mouth, manipulate objects, build structures, and tend to their eggs (Altman, 1971; Packard, 1988; Grasso, 2008; Hanlon and Messenger, 2018; Bidel et al., 2022; Buresch et al., 2022). In support of these behaviors, and like the arms themselves, each sucker can be moved independently and in near infinite degrees of freedom (Kier and Smith, 2002; Röckner et al., 2023). The suckers are additionally profound sensory structures, each containing an intricate sensory epithelium enriched with chemotactile and mechanoreceptors, allowing sensory detection of the environment in the absence of vision (Wells, 1964; van Giesen et al., 2020; Buresch et al., 2022; Kang et al., 2023).

The octopus arm houses a substantial nervous system, much of it dedicated to the control of the suckers (Young, 1963; Graziadei, 1971; Neacsu and Crook, 2024; Olson et al., 2025). There is a central nerve center: the massive axial nerve cord (ANC), equivalent to a spinal cord, running down the center of each arm. Within the ANC, a neuronal cell body layer (CBL) wraps around the neuropil (NP) forming a horseshoe pattern on the oral, or sucker, side (Graziadei, 1971; Olson et al., 2025). On the aboral side, or away from the sucker, there is a cerebro-brachial tract, comprising a pair of fiber tracts connecting the arms and the brain (Graziadei, 1971; Zullo et al., 2019). The CBL and NP of the ANC can be further divided into an aboral brachial territory with connections to the arm, and an oral sucker territory with connections to the sucker (Rowell, 1963; Olson et al., 2025). Down the long axis of the arm, this oral sucker territory of the ANC forms local enlargements over each sucker (Rossi and Graziadei, 1954; Olson et al., 2025; these enlargements are sometimes referred to as ganglia, see: Young, 1971; Nixon and Young, 2003; Nödl et al., 2016; Neacsu and Crook, 2024). From these enlargements, the ANC issues nerves fibers that spatially tile each sucker to create a topographic map (Olson et al., 2025).

The arm also harbors a peripheral nerve center for the sucker: the sucker ganglion (Graziadei, 1971). Given the strong coupling between the ANC and each sucker, what does the sucker ganglion do? Some early authors postulated a primarily motor role, with the sucker ganglion modulating local reflexes for the sucker stalk (Guérin, 1908). Others included a sensory role, with some sucker ganglion cell bodies being primary sensory cells (Rowell, 1963; Graziadei and Gagne, 1976). Lastly, others proposed a dual sensory and motor role (Martoja and May, 1955; Graziadei, 1965, 1971). Here, we investigated the role of the sucker ganglion by interrogating its molecular composition and circuitry with modern molecular and cellular methods.

## Materials and Methods

### Animals

Adult *Octopus bimaculoides* (California two-spotted octopus) were wild caught and purchased from the field collection venture, Aquatic Research Consultants, operated by Dr. Chuck Winkler (San Pedro, CA). After overnight FedEx delivery, octopuses were housed individually in 20-gallon artificial seawater (ASW) tanks, equipped with a carbon particle filter, an aquarium bubbler, and a UV light. Octopuses were fed daily with fiddler crabs, bivalves, or frozen shrimp. ASW was prepared by resuspending pharmaceutically pure sea salt (Tropic Marin “classic,” Wartenberg, Germany) in deionized water to a concentration of 33 g/liter.

Adults were anesthetized in 4% ethanol/ASW (EtOH/ASW, n = 13) and trans-orbitally perfused with 4% paraformaldehyde/phosphate-buffered saline solution (PFA/PBS, pH 7.4) delivered via a peristaltic pump (Cole Parmer Masterflex) through a 21½ gauge needle (Becton Dickinson). Throughout the procedure, hematopoietic tissue, consisting of left and right white bodies, was targeted alternately and iteratively. Arms were stored in 4% PFA/PBS overnight at 4°C. These arms were then either cut into 2-4 cm pieces for immediate processing or stored whole in diethyl pyrocarbonate (DEPC)-treated-PBS at 4°C.

These cephalopod experiments were performed in compliance with the EU Directive 2010/63/EU guidelines on cephalopod use, the University of Chicago Animal Resources Center oversight, and the University of Chicago Institutional Animal Care and Use Committees (Fiorito et al., 2015; Lopes et al., 2017).

### Tissue for Sections

Arm blocks were saturated with 30% sucrose/4%PFA/PBS at 4°C, rinsed with 30% sucrose/PBS, and infiltrated with 10% gelatin/30% sucrose/PBS for 1 hour at 50°C. Tissue was embedded in 10% gelatin/30% sucrose/PBS, post-fixed in 30% sucrose/4%PFA/PBS, and stored at -80°C. Serial arm sections were cut on a freezing microtome (Leica SM2000R) at 28-50-µm thickness and collected in DEPC-PBS. Sections were mounted and dried on charged, hydrophilic glass slides (TruBond380, Newcomer Supply, Middleton, WI) before storage at -80°C until further processing.

### cDNA Synthesis and Cloning

For RNA extraction, dissected octopus central brain and arm tissues were flash-frozen on dry ice and stored at -80°C. After homogenizing tissue with a micropestle, RNA was extracted with Trizol Reagent (Invitrogen) and phasemaker tubes (Invitrogen). cDNA was synthesized using the SuperScript III 1st-strand cDNA kit (Invitrogen) following manufacturer instructions. RNase-free water (Sigma-Aldrich) was used to dilute cDNA, which was stored at -20°C until use.

PCR primers were designed with MacVector software (version 12.6.0) or PrimerBlast from NCBI (Table 1). PCR reaction solutions were incubated using the T100 thermocycler from BioRad at 95°C for 5 minutes. Then, the solutions underwent 35-40 rounds of amplification cycles: 95°C for 30 seconds, 52-57°C for 45 seconds, and 72°C for 1 minute. A final elongation step was performed at 72°C for 10 minutes.

**Table 1:**
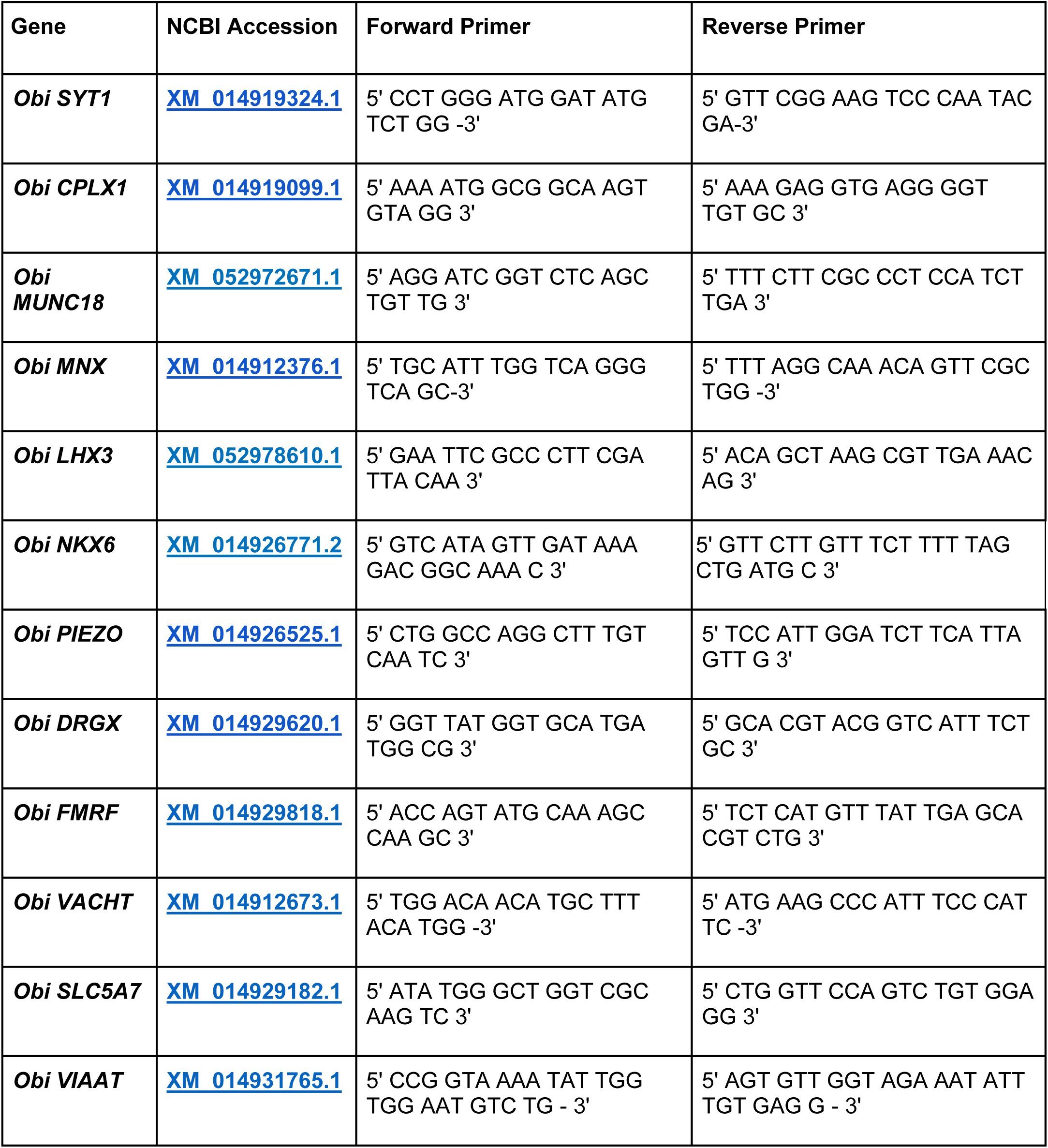
Transcripts amplified for in situ hybridization.

The sequences for the neurotransmitter genes *GAD* and *CHAT* were cloned by Dr. Rahul Parnaik using degenerate PCR. The sequence for *GAD* cDNA matches 18-364 of XM_014922646. That for *CHAT* matches 1085-2063 of XM_014912671.

PCR reaction products were ligated into a pGEM T-Easy plasmid (Promega), and closed inserts were Sanger sequenced by the University of Chicago Comprehensive Cancer Center DNA Sequencing Facility. To generate antisense templates, plasmids were linearized with SacII (New England Biolabs, Cat#: 50812058) or SpeI (New England Biolabs, Cat#: 50811989). Phenol-chloroform extraction was performed on the template. Antisense digoxigenin (DIG)-labeled riboprobes (Sigma-Aldrich, Cat#: 11277073910) were transcribed with SP6 or T7 RNA polymerase (New England Biolabs). The residual template was digested at 37°C for 15 minutes with RNase-free DNase I (Sigma Aldrich). Riboprobes were ethanol-precipitated and stored in 100μl of DEPC-H_2_O at -20°C until use.

### In situ hybridization (ISH)

Slides of sectioned tissue were equilibrated to room temperature, post-fixed in mailers in 4% PFA/PBS, then washed in DEPC-PBS. Following a 15-minute incubation at 37°C in proteinase K solution (Sigma-Aldrich; 1.45 μg proteinase K per milliliter of 100 mM Tris-HCl [pH 8.0], 50 mM EDTA [pH 8.0]), slides were post-fixed for 15 minutes in 4% PFA/PBS, rinsed in DEPC-PBS, and acclimated to 72°C for 1 hour in hybridization solution (50% formamide, 5x SSC, 1% SDS, 200 μg/ml heparin, 500 μg /ml yeast RNA). Slides were transferred to mailers with 1-2 mg antisense riboprobe in 15 mL hybridization solution and incubated overnight at 72°C. The next day, slides were treated in preheated Solution X (50% formamide, 5x SSC, 1% SDS) at 72°C, washed in room temperature TBST (Tris-buffered saline with 1% Tween 20), and blocked at room temperature for one hour in 10% DIG buffer (Roche) in TBST. Meanwhile, anti-DIG Fab fragments conjugated to alkaline phosphatase (Sigma-Aldrich, Cat#: 11093274910, RRID: AB_514497) were preabsorbed with octopus embryo powder in 1% DIG buffer in TBST for at least 1 hour. Slides were incubated on a rocker overnight at 4°C in preabsorbed antibody diluted to a final concentration of 1:5000 in 10% DIG buffer in TBST.

The next day, slides were washed in TBST and moved into freshly prepared NTMT (100 mM Tris-HCl [pH 9.5], 100 mM NaCl, 50 mM MgCl2, 1% Tween 20) for 10 minutes. To demonstrate alkaline phosphatase activity, the slides were incubated in NTMT containing nitro blue tetrazolium (NBT, 100mg/mL in 70% dimethyl formamide/30% DEPC-H20, Gold Biotechnology, St. Louis, MO) and 5-bromo-4-chloro-3-indolyl phosphate (BCIP, 50mg/mL in 100% dimethyl formamide, Gold Biotechnology, St. Louis, MO). Color reaction was monitored for up to 5 days. After such time, slides were washed in TBST overnight, dehydrated through an ethanol series, cleared in Histoclear (National Diagnostics, Atlanta, GA), and coverslipped with Eukitt (Sigma-Aldrich).

In situ hybridization with no probe and a mouse probe were used as controls. No staining was observed in the no probe condition. The heterologous probe elicited labeling restricted to the edge of the sucker, as illustrated in Fig. 1 b, c, and f.

**Figure 1:**
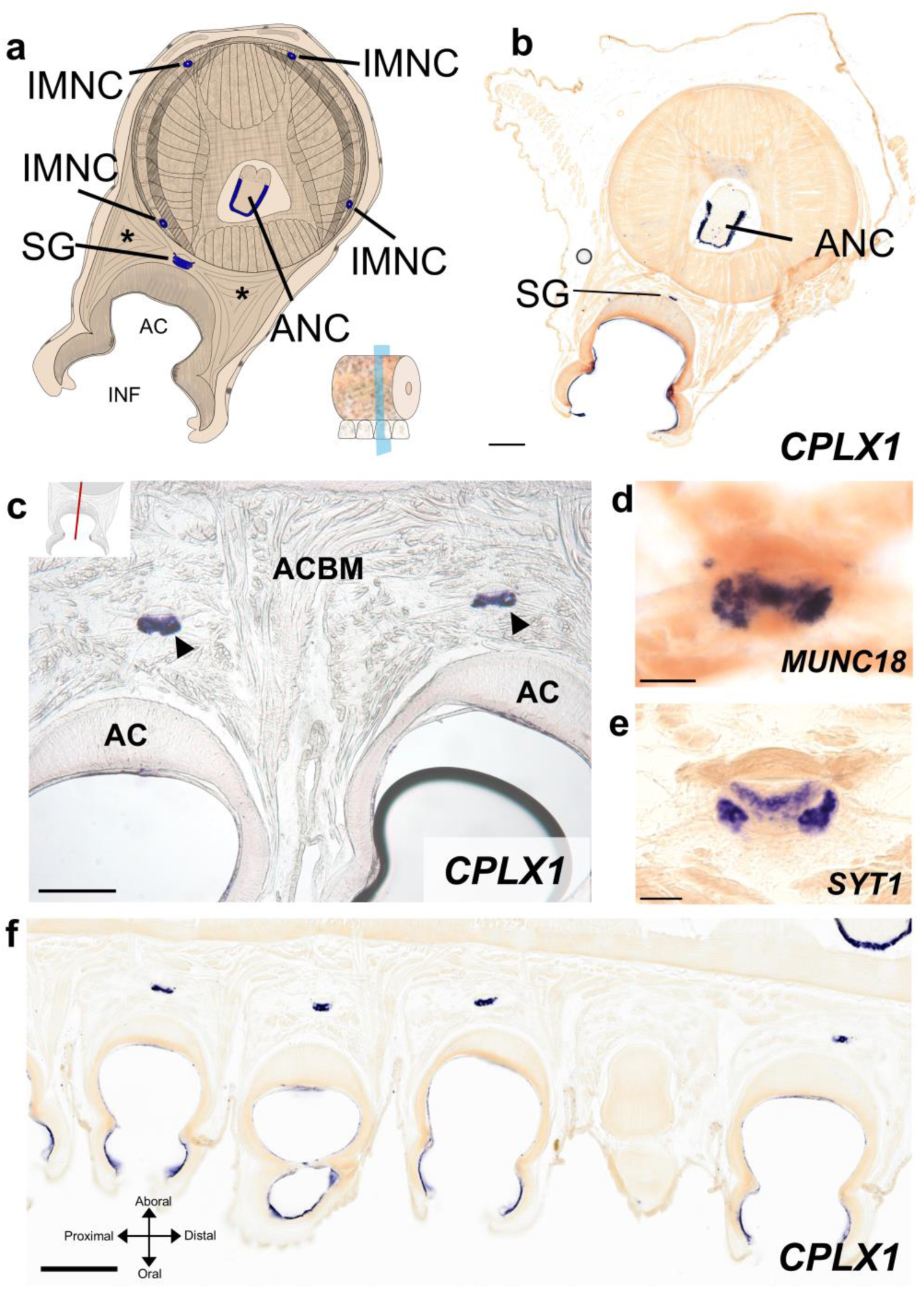
Geography of octopus arm and location of sucker ganglion. **a,** Diagram of a transverse cross section of octopus arm. The major components of the arm nervous system are highlighted in blue. Key indicates transverse plane of section. *denotes acetabulo-brachial muscles (ACBM) **b,** Transverse arm section with in situ hybridization (ISH) for the neuronal gene complexin (*CPLX1*). Neuronal cell bodies in the axial nerve cord (ANC) and sucker ganglion (SG) are labeled. Scale bar: 100 μm **c-f,** Longitudinal sections through the octopus arm, as indicated by inset in c. In the longitudinal plane, the coordinate axes correspond to aboral-oral, proximal-distal, as indicated in f. **c,** ISH of *CPLX1* on a longitudinal section of octopus arm. There is an SG embedded in the ACBM for every sucker. Arrowheads point to the SG. Scale bar: 250 μm **d-e,** The SG is enriched with other panneuronal markers. **d,** *MUNC18.* **e,** *SYT1.* Octopus have many isoforms of *SYT*, so *SYT1* may not label all neurons. Scale bars: 50 μm. **f,** ISH of *CPLX1* on a longitudinal section of octopus arm across many suckers. The location of the SG varies based on the location of the sucker. Scale bar: 500 μm. AC, acetabulum; ACBM, acetabulo-brachial muscles; ANC, axial nerve cord; IMNC, intramuscular nerve cord; INF, infundibulum; SG, sucker ganglion.

### Hematoxylin and Eosin (H&E) Staining

Slides were incubated in DEPC-PBS for 4-6 hours at 72°C to melt off gelatin. Then, slides were rehydrated in room temperature deionized water, incubated in Mayer’s Hematoxylin (ScyTek, Cat#: HMM500) for 1 minute, and rapidly rinsed in water to prevent overstaining. After a 15 second incubation in a bluing solution (ScyTek, Cat#: BRT500), slices were dehydrated in an ethanol series (70%, 95% and 100%) and placed in Eosin Y (ScyTek, Cat#: EYB500) for 1 minute. Slides were rinsed in 100% EtOH, cleared in Histoclear, and mounted with Eukitt.

### Picrosirius Red

The presence of connective tissue was examined with Picrosirius Red (Abcam, Cat#: ab150681). De-gelatinized slides were rinsed 3 times with deionized water then rehydrated for 1-minute in deionized water. Sections were incubated in Picrosirius Red solution for 5 minutes, de-stained with 2 quick rinses of acetic acid wash, dehydrated in 100% EtOH, and mounted with Eukitt.

### Immunohistochemistry

Mouse monoclonal antibody 6–11B-1 (1:500 dilution of ascites fluid; RRID: AB_477585, Sigma-Aldrich, Cat#: T6793) and mouse monoclonal antibody SMI-31 (1:500 dilution; RRID: AB_2564641, BioLegend, Cat# 801601) were used to label neuronal processes in the octopus arm. Clone 6–11B-1, which was isolated following immunization with sea urchin sperm flagella protein preparations, has been employed to label axon tracts in vertebrate and invertebrate nervous systems (Piperno, Gianni and Fuller, Margaret T., 1985; Chitnis and Kuwada, 1990; Shigeno and Yamamoto, 2002; Baratte and Bonnaud, 2009). It recognizes an acetylated α-tubulin (acTUBA) epitope found broadly but not universally across microtubules (Piperno, Gianni and Fuller, Margaret T., 1985; LeDizet and Piperno, 1991). Clone SMI-31 recognizes a phosphorylated epitope on neurofilament in mammals and neurofilament 220 in squid (Sternberger and Sternberger, 1983; Grant et al., 1995; Grant and Pant, 2016). Alexa Fluor® 488 AffiniPure Goat Anti-Mouse IgG (Jackson ImmunoResearch, Cat#: 115-545-003, RRID: AB_2338840) and Cy™3 AffiniPure Donkey Anti-Mouse IgG (Jackson ImmunoResearch, Cat#: 715-165-151, RRID: AB_2315777) were utilized as secondaries (1:500 dilutions).

For section immunohistochemistry, slides were washed in DEPC-PBS containing 1% Tween 20 (PBST), incubated for 30 minutes in a proteinase K solution at 37°C (Sigma-Aldrich, Cat#: 03115828001; 19.4 μg proteinase K per milliliter of PBST), then returned to room temperature with a post-fix in 4% PFA/PBS for 15 minutes. Following washes in PBST, slides were blocked in 10% goat serum/PBST (Fisher Scientific, Cat#: 16210072) for 1 hour and incubated in primary antibody diluted in 1% goat serum/PBST for 4 days at 4°C. Slides were washed in PBST for 3x30 minutes. Then, sections were incubated for 2 days at 4°C with secondary antibody along with 0.01 mg/ml 4′-6-diamidino-2-phenylindole (DAPI; Sigma-Aldrich, Cat#: D9542) to label cell nuclei fluorescently and either 0.2 µl/ml phalloidin-iFluor 594 (Abcam, Cat#: ab176757) or phalloidin-iFluor 488 (Abcam, Cat#: ab176753) to label filamentous actin (F-actin). Slides were then washed with PBST and cover slipped with Fluoromount G (Southern Biotech, Birmingham, AL).

For whole mount immunohistochemistry, 0.5 cm – 1 cm transverse slices and dissected suckers were washed in PBST, dehydrated in a graded methanol series (25%, 50%, 75% in PBST), and stored overnight at -20°C. Slices were rehydrated the next day in a graded methanol series (75%, 50%, 25% in PBST), rinsed in PBST, incubated for 1 hour at 37°C in a proteinase K solution (19.4 μg proteinase K per milliliter of PBST), and post-fixed for 15 minutes in 4% PFA/PBS. Following washes in PBST, tissue was blocked for 1 hour in 10% goat serum/PBST and incubated in primary antibody for 7 days at 37°C. Following a day of PBST washes, tissue was transferred to secondary antibody for 7 days at 4°C. Tissue was rinsed with PBST, washed with DEPC-PBS, and post-fixed for 4 days at 4°C. Blocks were rinsed in DEPC-PBS and then cleared in a modified version of FRUIT with incubations in 35%, 40%, 60%, 80%, 100% FRUIT, each for 24 hours (Hou et al., 2015). Until imaging, slices were stored in 100% FRUIT at 4°C.

### Imaging and image analysis

Lightfield and fluorescent-labeled tissue was studied with a Zeiss Axioskop 50 upright microscope and a Leica MZ FLIII stereomicroscope, both outfitted with the Zeiss AxioCam digital camera and AxioVision 4.5 software system. Selected sections were also studied on a Leica SP5 Tandem Scanner Spectral 2-photon confocal microscope (Leica Microsystems, Inc., Buffalo Grove, IL) or scanned by an Olympus VS200 Research Slide Scanner (Olympus / Evident, Center Valley, PA) with a Hamamatsu ORca-Fusion camera (Hamamatsu Photonics, Skokie, IL). Whole mounts were imaged on a LaVision BioTec UltraMicroscope II (Miltenyi Biotec, Bergish Gladbach, Germany) run by ImSpector Pro v. 7_124 software (LaVision BioTec, Bielefeld, Germany). Collected images were corrected for contrast and brightness and false colored in FIJI (version 2.1.0/1.53c; National Institutes of Health (NIH)).

### Proximal-distal analysis

Two series were created to examine sucker ganglia size changes along the proximal-distal axis of the arm from the edge of crown webbing to tip. The first series consisted of 3 evenly spaced arm blocks from an arm on the right side (arm R1) of one animal sectioned longitudinally and stained with acTUBA. The second series consisted of 3 evenly spaced blocks from an arm on the left side (arm L3) of a second animal sectioned longitudinally and stained with H&E. The longitudinal width of each sucker ganglion along the proximal-distal axis was manually measured at the widest point using the line tool in FIJI (Fig. 2a, b). The longitudinal widths of the sucker ganglion were then compared to the longitudinal widths of the corresponding suckers (SG width/sucker width). Sucker measurements are reported in Olson et al. (2025).

**Figure 2:**
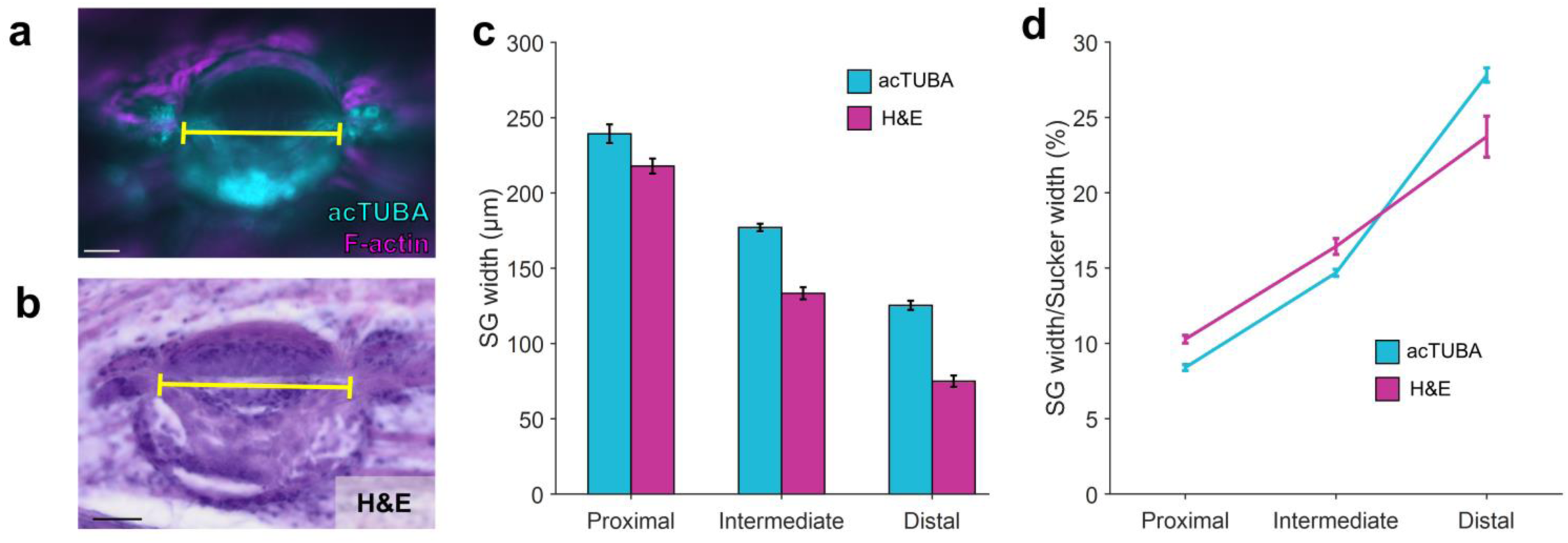
Sucker ganglion width changes along the proximal-distal axis. **a-b,** Example longitudinal sections through sucker ganglia (SG) used for measurements. Yellow line indicates location of measurement. Scale bars: 50 μm. **a,** SG stained with acetylated alpha tubulin (acTUBA, cyan) and F-actin (magenta). **b,** SG stained with hematoxylin and eosin (H&E). **c,** SG width decreases down the proximal (P)- distal (D) axis, error bars +/- sem. **d,** The width of the SG relative to sucker width increases along the P-D axis, error bars +/- sem. acTUBA, acetylated alpha tubulin; H&E, hematoxylin and eosin; SG, sucker ganglion.

### Nerve analysis

Large scale nerve analysis was facilitated by segmenting 2 transversely imaged whole mount sucker ganglia, immunostained with acTUBA and SMI-31, and 2 horizontally imaged whole mount sucker ganglia, immunostained with acTUBA. Nerve fibers connecting to the sucker ganglion were traced using Simple Neurite Tracer (SNT) in FIJI (Arshadi et al., 2021).

### Nerve fascicle diameters

Using the path fitting feature in SNT, the radii of each node of the paths of the 2 horizontally imaged sucker ganglion were found (Arshadi et al., 2021). Then, using the SNT API and a python script, the radii of the nodes corresponding to the first 300 μm of each path were extracted, smoothed and averaged. The average radius for each path was multiplied by 2 to determine the diameter of the nerve fascicle. The distribution of nerve fascicle diameters was plotted as a histogram using Matplotlib (v. 3. 7.3).

### Nerve root distribution

The SNT API was used to find the 3D coordinates of the roots of the traces from the 2 transversely imaged sucker ganglia and from 1 horizontally imaged sucker ganglion (Arshadi et al., 2021). The coordinates of roots of the nerves were translated and rotated so that the center of all the roots was set as the origin. Subsequently, each root coordinate was normalized so that the vector from the origin to the tip had a magnitude of 1, creating a circle. The spatial distribution of roots from the 3 sucker ganglia was realized in the aboral-oral and external-internal plane with a colormap corresponding to the location of the root in that plane using Matplotlib. The spatial distribution of roots from the horizontally imaged sucker ganglion was also realized with a colormap corresponding to the diameter of the nerve fiber.

### Nerve targets

The 3D spatial coordinates, roots, and tips of the nerves from the horizontally imaged and transversely imaged sucker ganglia, both stained with acTUBA, were found using the SNT API (Arshadi et al., 2021). The average trajectory of each nerve was computed by creating a vector from the root of the nerve to the center of the process (Fig. 7i). The unit vector for each average trajectory was found. The distribution of the average trajectories for the transversely imaged sucker ganglion was realized in the aboral-oral and external-internal plane with a colormap corresponding to nerve root location using Matplotlib. The distribution of the average trajectories for the horizontally imaged sucker ganglion was realized in the proximal-distal and external-internal plane with a colormap corresponding to nerve target location in the plane.

## Results

We confirmed the presence and position of the sucker ganglion in *O. bimaculoides* with in situ hybridization (ISH) for pan-neuronal markers (Fig. 1b, c, d, e). As in other species, the sucker ganglion is embedded in the acetabulo-brachial muscles (ACBM) comprising the sucker stalk aboral, or deep, to the acetabulum of each sucker (Fig. 1b, f). Because of its position within the sucker musculature, the sucker ganglion is a mobile structure. As demonstrated in Fig. 1f, the sucker ganglion moves with the sucker as the sucker muscles contract.

Suckers decrease in width along the proximal-distal axis of the arm. The sucker ganglion likewise decreases in width, though it maintains its ellipsoid shape down the long axis of the arm (Fig. 2c). We asked if this decrease is isometric with the decrease in sucker width. We found instead an allometric relationship: ganglia from the distal region of the arm were larger relative to the width of their respective sucker than ganglia found in the intermediate or proximal regions (Fig. 2d). This difference could be due to a developmental constraint on the size of the circuit or underlying functional differences across suckers located at different arm positions.

### Internal composition of the sucker ganglion

The octopus sucker ganglion has an unusual structure (Graziadei, 1971; Supplemental Video 1). Instead of neuronal cell bodies surrounding a central neuropil (Richter et al., 2010), the sucker ganglion neuropil (NP) is predominantly arranged in a cap on the aboral surface (acTUBA labeling; Fig. 3d). A dense cell body region is located just orally of this NP cap, that is, a step closer to the sucker (Fig. 3e, f, h-k). This cell body region is composed of small cell bodies situated in a basket of larger cell bodies (Fig. 3e, f). At the junction of the NP cap and cell body region is an inner belt (Fig. 3d, e). This inner belt is composed of circumferentially running fiber bundles with large cell bodies lining its outer edge (Fig. 3d, e, h-k). In addition, connective tissue identified with Picrosirius Red staining forms a distinct plate between the NP cap and cell body region, separating these two territories (Fig. 4). On the oral side of the sucker ganglion, that is closest to the sucker, there is a small pocket of neuropil and an additional belt of circumferential fibers (“outer belt”; Fig. 3f, g). Unlike the inner belt, however, the outer belt appears devoid of neuronal cell bodies (Fig. 3f, g).

**Figure 3:**
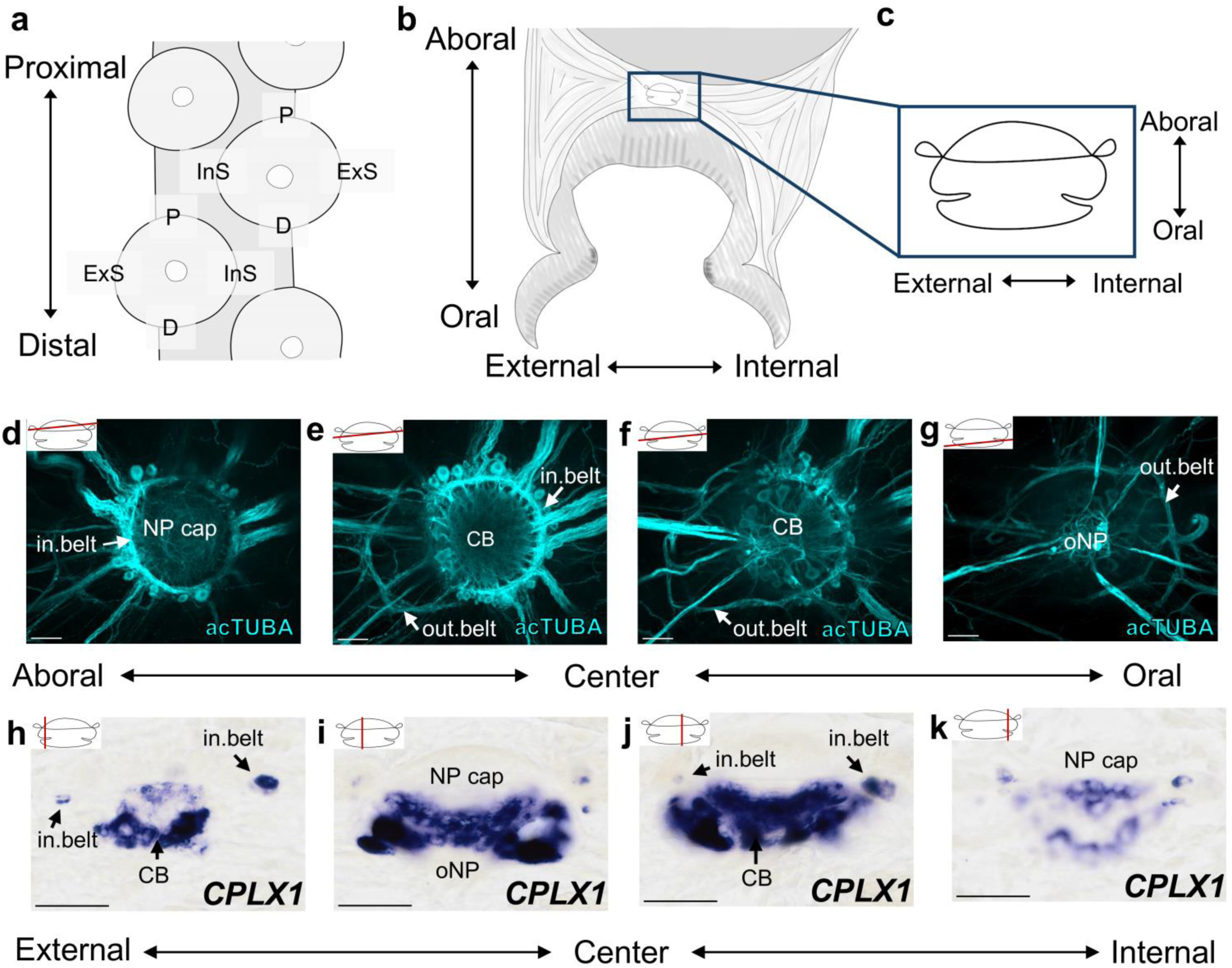
Internal neural organization of the sucker ganglion. **a-c,** The axes for a sucker. **a,** An en face, horizontal cartoon of the suckers showing that the suckers are arranged in two offset rows. In the horizontal plane, each sucker has a proximal (P)- distal (D) axis, and an internal side (InS)- external side (ExS) axis. The P-D axis corresponds to the P-D axis for the arm as whole (that is, from head to arm tip). The InS corresponds to the side of the sucker that hugs the midline of the arm. The ExS corresponds to the side of the sucker that is situated on the outer edge of the arm. **b,** A transverse cartoon of a sucker. In the transverse plane, each sucker has an aboral-oral and ExS-InS axis. The aboral side is the side further from the sucker, the oral side is the side closest to the sucker. **c,** A cartoon of a transverse section through the sucker ganglion (SG). Like the whole sucker, the SG has an aboral-oral axis and an ExS-InS axis in the transverse plane. **d-g,** Organization of the SG demonstrated by acetylated alpha tubulin (acTUBA, cyan) immunostaining. Horizontal virtual slices produced by fusing multiple 1 μm confocal sections from aboral (d) to oral (g). See Supplemental Video 1 for a 3-D rendering of this dataset. **d,** Most aborally is a neuropil (NP) cap and an internal belt (in.belt) that is composed of circumferential fibers and large cell bodies. **e,** The in.belt is situated aboral to the cell body (CB) region. **f,** Oral to the CB territory is an outer belt (out.belt), composed of circumferentially running fiber bundles. **g,** On the oral side of the SG is a pocket of oral NP (oNP). Inset highlights the slight asymmetry of the horizontal sections through the SG. **h-k,** Longitudinal sections from external (h) to internal (k) through the SG prepared for in situ hybridization with *CPLX1* demonstrating the location of neuronal cell bodies. Neurons are located in the in.belt and are dense throughout the full extent of CB region. Scale bars: 50 μm. acTUBA, acetylated alpha tubulin; CB, cell bodies; D, distal; ExS, external side; in.belt, internal belt; InS, internal side; NP cap, neuropil cap; oNP, oral neuropil; out.belt, outer belt; P, proximal.

**Figure 4:**
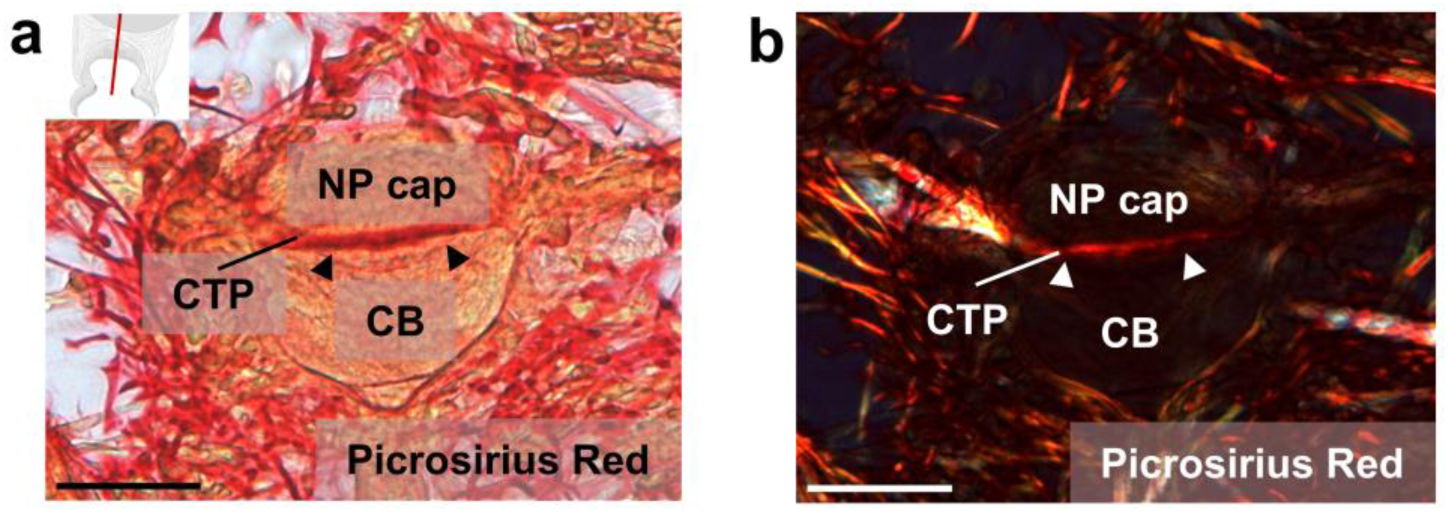
Connective tissue splits the sucker ganglion. **a-b,** Picrosirius Red staining of a longitudinal section through the sucker ganglion (SG). Red denotes the presence of connective tissue. **a,** Brightfield image depicting the full structure of the SG. Black arrowheads point to connective tissue plate (CTP) between the cell body (CB) region and neuropil (NP) cap. **b,** Polarized light image of the same field, highlighting the red labeling of connective tissue. White arrowheads point to CTP. Scale bars: 50 μm. CB, cell bodies; CTP, connective tissue plate; NP cap, neuropil cap.

### Molecular characterization of the sucker ganglion cell bodies

With ISH, we investigated the molecular composition of the cell body region of the sucker ganglion, first examining sensory and motor neuron markers (Jessell, 2000; Coste et al., 2010; Nomaksteinsky et al., 2013). We found positive expression for two motor neuron makers, *MNX* and *LHX3*, and the mechanosensitive ion channel marker *PIEZO* (Fig. 5a, b, c). *PIEZO* expression is found in scattered cells throughout the cell body territory, whereas *MNX* and *LHX3* labeling lies at its lateral edges and in its center. *PIEZO* expression was also present in a subset of cells in the inner belt (Fig. 5a). Interestingly, we failed to detect expression for *DRGX*, the dorsal root ganglion homeobox gene, and *NKX6,* a transcription factor marking motor neurons, despite positive labeling for both of these markers in the ANC (Fig. 5g, h; Olson et al., 2025). Hence, while the sucker ganglion contains expression for both sensory and motor markers, its molecular profile differs from that of the ANC.

**Figure 5:**
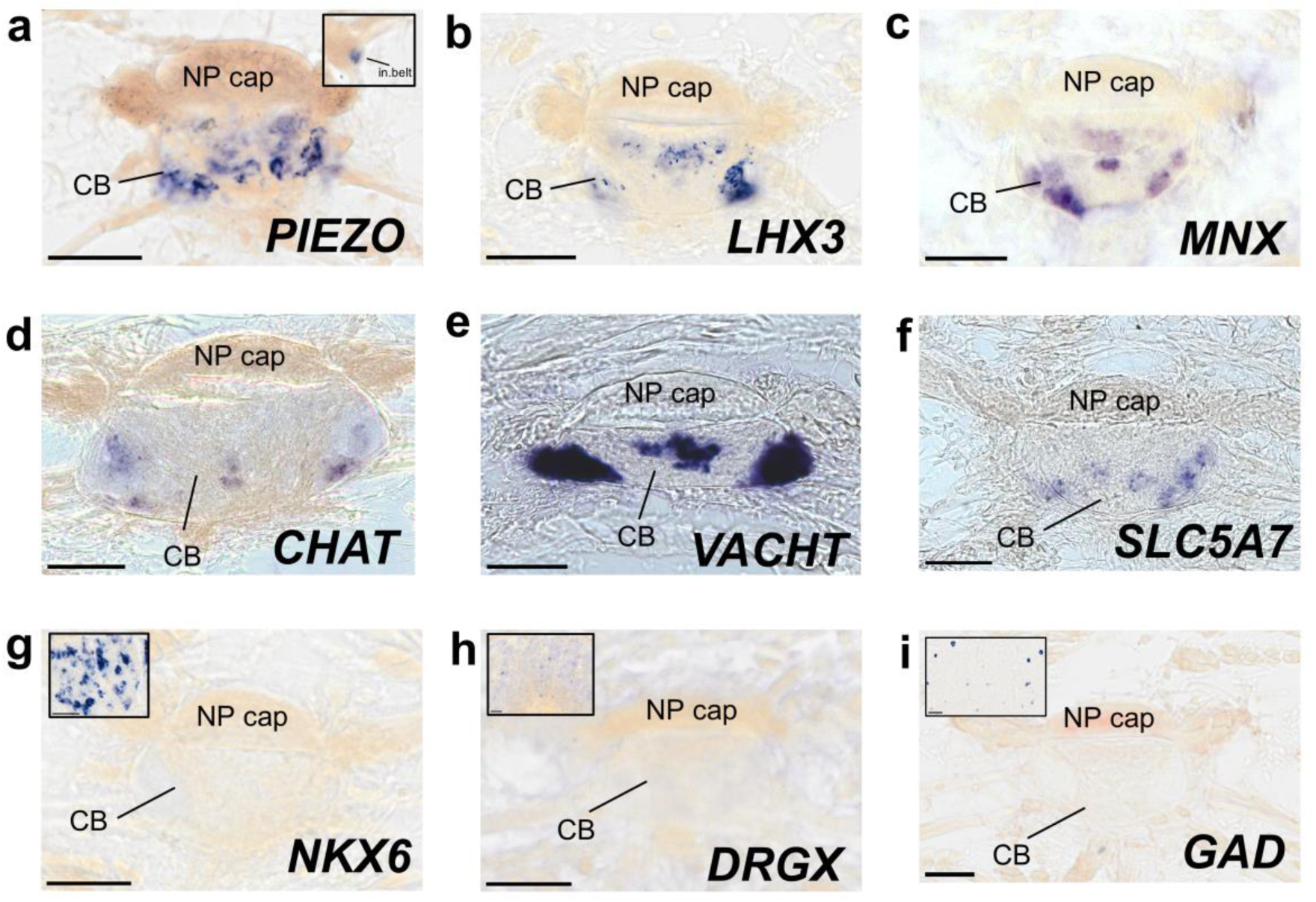
Gene expression patterns in the cell body territory of the sucker ganglion. **a-c,** the sucker ganglion (SG) is enriched with the sensory and motor neuron markers, **a,** In situ hybridization (ISH) for the mechanosensory ion channel marker *PIEZO*. Inset demonstrates *PIEZO* labeling in the inner belt (in.belt), **b,** ISH for LIM homeobox 3 transcription factor (*LHX3*), a motor neuron marker, **c,** ISH for motor neuron and pancreas homeobox transcription factor (*MNX*). **d-f,** Presysnaptic markers for cholinergic neurons label a subset of neurons in the cell body region. **d,** ISH for choline acetyl transferase (*CHAT*), **e,** ISH for vesicular acetylcholine transporter (*VACHT*) and **f,** ISH for solute carrier family 5 member 7 (*SLC5A7*). **g-h,** The molecular profile of the SG differs from that of the ANC. No positive labeling was detected in the SG for **g,** NK6 homeobox transcription factor (*NXK6*), a motor neuron marker, **h,** Dorsal root ganglion homeobox transcription factor (*DRGX*), **i,** glutamate acid decarboxylase (*GAD*), despite positive expression in the axial nerve cord (insets in **g, h, i**). All sections longitudinal. CB, cell bodies; in.belt, inner belt; NP cap, neuropil cap. Scale bars: 50 μm.

Graziadei (1965) reported the existence of muscle receptors in the ACBM around the sucker ganglion. With acTUBA immunostaining, we identified many cells embedded in the ACBM which issue processes to nerve fascicles connected to the sucker ganglion (Fig. 6). Some of these cells are multipolar with additional processes extending into the local musculature (Fig. 6b, c). Some cells cluster together in groups (Fig. 6d). Others are located close to the sucker ganglion itself (Fig. 6e). We also found that the ACBM are enriched with expression of *PIEZO* and *DRGX*, indicating that these cells may be sensory in nature. This suggests the sucker ganglion handles sensory input from cells within the ganglion itself as well as from the local musculature.

**Figure 6:**
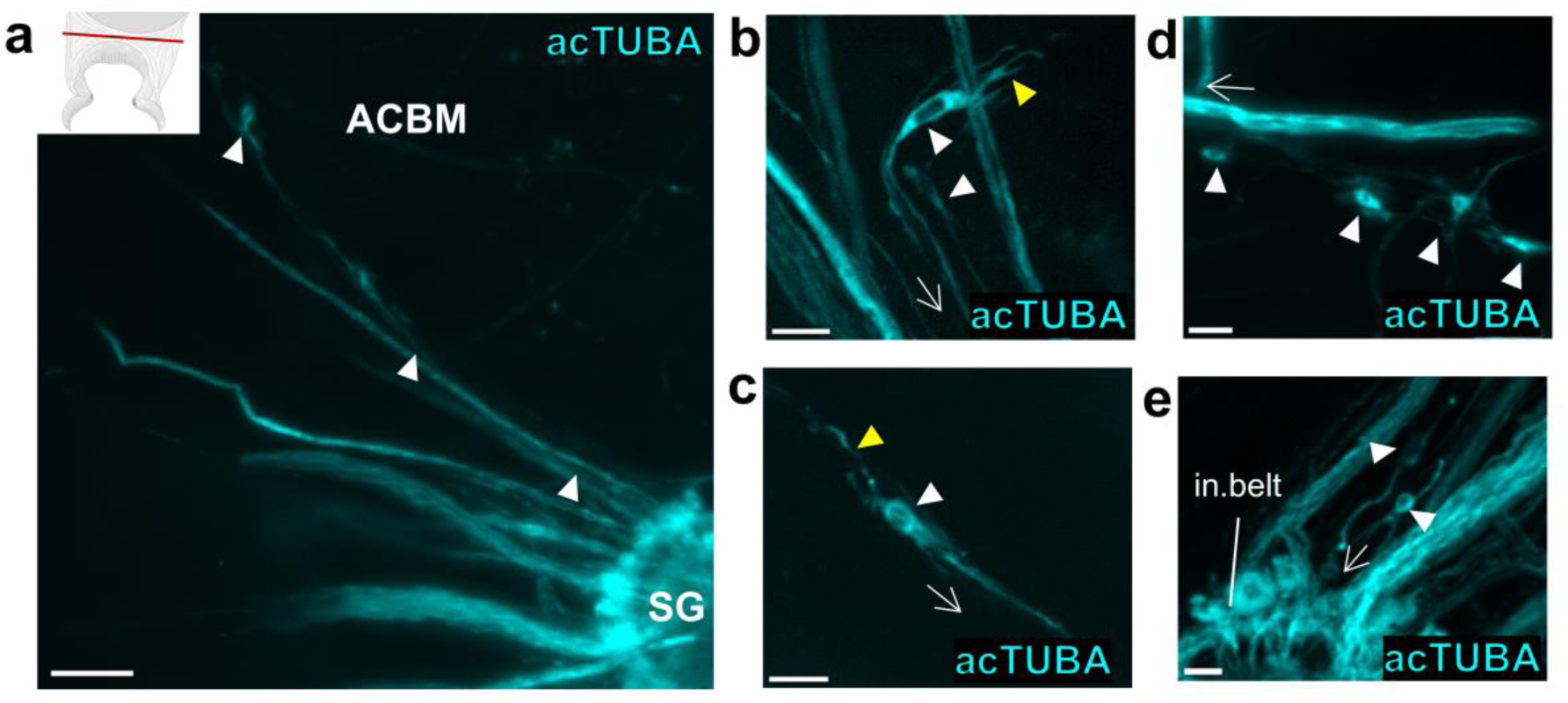
Cells in the local musculature feed back to the sucker ganglion. **a,** Maximum projection of the sucker ganglion (SG) and surrounding musculature in the horizontal plane stained with acetylated alpha tubulin (acTUBA, cyan). Arrowheads point to cell bodies in the musculature that feeds back to SG. Scale bar: 100 μm. **b-e,** Close-up images of cells in musculature stained with acTUBA. **b-c,** Cells exhibit a complex morphology. Yellow arrowheads point to processes that extend into the musculature, white arrowheads indicate cell bodies. Arrows point in direction of the SG. Scale bars: 25 μm. **d,** Some cells are located in clusters. White arrowheads indicate cell bodies. Arrow points in direction of SG. Scale bar: 50 μm. **e,** Some cells are located close to the SG. White arrowheads indicate cell bodies. Arrow points in direction of the SG. Scale bar: 25 μm. ACBM, acetabulo-brachial muscles; acTUBA, acetylated alpha tubulin; in.belt, inner belt; SG, sucker ganglion.

We interrogated the sucker ganglion with ISH for other neural subtype markers. The sucker ganglion is enriched for expression of presynaptic cholinergic neuron markers (*VACHT, SLC5A7*, and *CHAT*). Unlike in the ANC, where we found expression for these cholinergic neuron markers throughout the entirety of the CBL, *VACHT, SLC5A7,* and *CHAT* expression in the sucker ganglion is found at the lateral edges of the ganglion and the center, similar to the labeling pattern of the motor neuron markers (Fig. 5d, e, f). Glutamate decarboxylase (*GAD*) and vesicular inhibitory amino acid transporter (*VIAAT*) serve as markers of fast GABAergic inhibitory neurons. We failed to detect significant expression for either of these inhibitory neuron markers in the sucker ganglion in experiments presenting positive labeling in the ANC (Fig. 5i). Rapid ganglionic inhibition therefore would need to be mediated by another neurotransmitter, such as acetylcholine, or issued from the ANC.

The expression pattern of *FMRF*, which encodes a neuromodulatory neuropeptide, was striking in the sucker ganglion. In the ANC, it is broadly and abundantly expressed (Fig. 7a; see also Winters-Bostwick et al., 2024). In the sucker ganglion, however, it is very restricted (Fig. 7b). Moreover, the distribution of labeling is asymmetric within the ganglion, with labelling found only on the external side, or the side that corresponds to the outer edge of the sucker (Fig. 7c, see Fig. 3a). This asymmetry in molecular identity could underlie an asymmetry in function.

**Figure 7:**
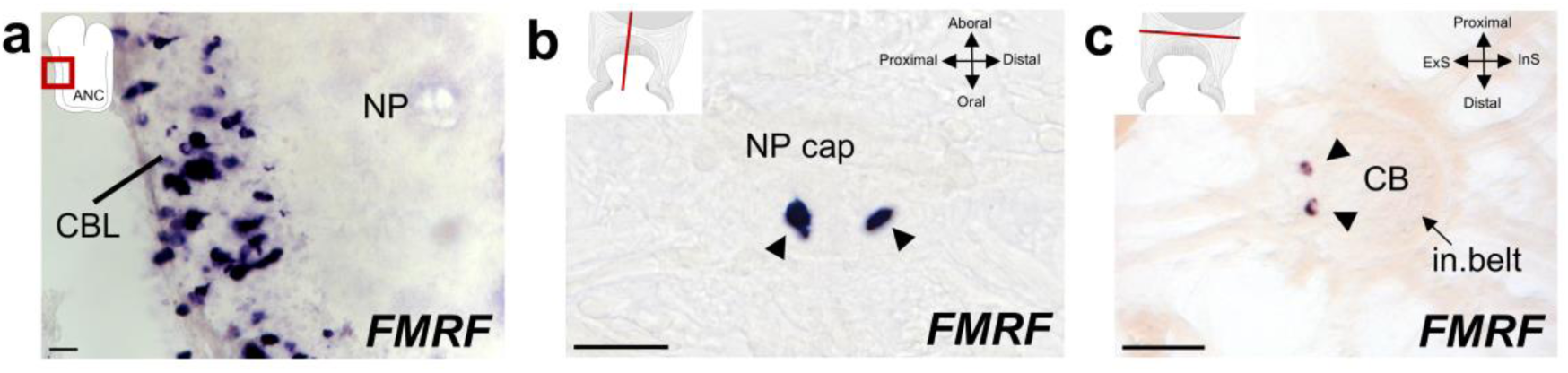
Asymmetric gene expression of *FMRF* in the sucker ganglion. **a,** Transverse section of the axial nerve cord with in situ hybridization (ISH) for *FMRF*. *FMRF* expression is found abundantly throughout the cell body layer. Inset indicates the location of the panel image **b,** Longitudinal section through the sucker ganglion (SG) with ISH for *FMRF*. Arrowheads point to *FMRF* labeling of a small subset of cells. **c,** Horizontal section through the SG with *FMRF* ISH. *FMRF* labeling is restricted to the external side of the SG, as indicated by arrowheads. ANC, axial nerve cord; CB, cell bodies; CBL, cell body layer; ExS, external side of sucker; InS, internal side of sucker; in.belt, inner belt; NP, neuropil; NP cap, neuropil cap. Scale bars: 50 μm.

### Nerve fiber organization

Many fiber fascicles issue from the sucker ganglion (Fig. 8a). These bundles vary in diameter from 6 µm to 45 µm, with the bulk falling in the 11-20 µm range (Fig. 8b). We examined the location of the fascicles around the perimeter of the sucker ganglion. We found that the roots are primarily distributed along the lateral sides and oral underside, forming a U-shape (Fig. 8c, d). There are notable gaps in this pattern, with a prominent gap corresponding to the NP cap, and two bilaterally symmetric gaps corresponding to a bulge in the cell body territory (Fig. 8c, d). The largest nerve bundles depart from the aboral portion of the ganglion (Fig 8e, f). This territory falls in between the aboral and lateral gaps (Fig. 8g). These results suggest a regionalization along the aboral-oral axis within the ganglion based on root location.

**Figure 8:**
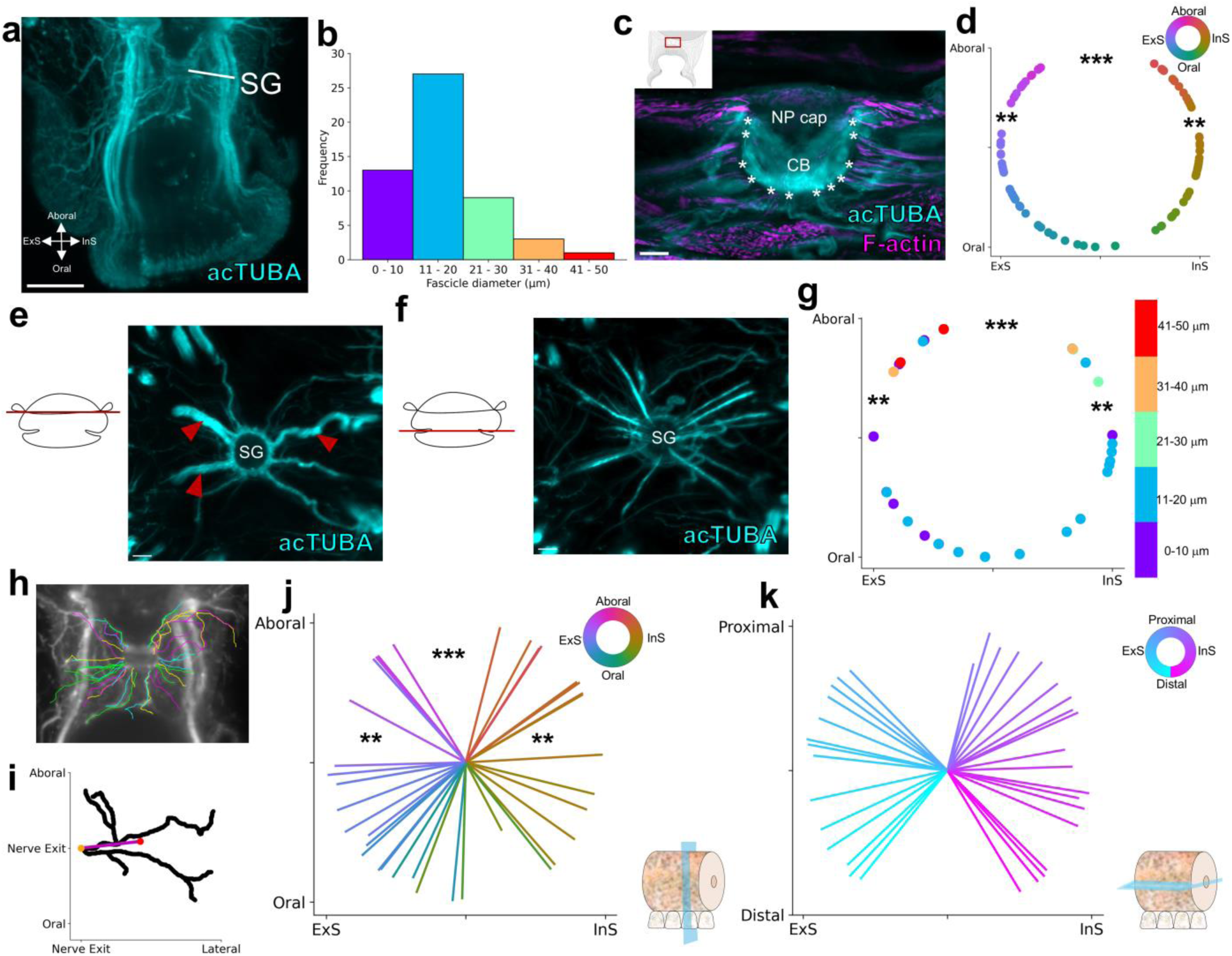
Distribution of sucker ganglion nerve fibers. **a,** Maximum projection of a whole mount of a sucker stained with acetylated alpha tubulin (acTUBA, cyan). Many fascicles enter and exit the sucker ganglion (SG). Scale bar: 250 μm. **b,** Distribution of fascicle diameters. Diameters range from 6-45 μm, with the peak in the 11-20 μm range. **c,** Transverse section through the SG stained with acTUBA (cyan) and F-actin (magenta). Location of nerve fiber exits are denoted with *. Scale bar: 50 μm. **d,** Distribution of fiber exits around the SG (n = 3 ganglia, 2 stained with acTUBA, 1 stained with SMI-31) plotted in the transverse plane. There is a gap corresponding to the neuropil (NP) cap (***) and two gaps corresponding to a protrusion of the cell body region (**). Color key by location of root in the aboral-oral, external-internal plane. **e-g,** The largest nerve fibers exit aborally. **e,** Horizontal maximum projection of the aboral portion of the SG stained with acTUBA. Red arrow heads indicate large nerve bundles. **f,** Horizontal maximum projection of the oral portion of the sucker ganglion stained with acTUBA. No large nerve bundles are present. Scale bars: 100 μm. **g,** Distribution of fiber exits around the SG (n = 1 ganglion, stained with acTUBA) plotted in the transverse plane. Color key corresponds to size of fascicles. The largest fiber bundles exit aborally, corresponding to the portion between the NP cap and cell body protrusion. **h,** Example of SG nerves segmented out of transverse whole mount slice stained with acTUBA. **i,** Schematic of the average trajectory, which is a vector created from the nerve exit point to the center of the processes. **j,** Distribution of average trajectories in the transverse plane (n = 1 ganglion, stained with acTUBA). Color key corresponds to nerve exit location in the aboral-oral, external-internal plane. Nerve targets correspond to nerve exit. Key illustrates the transverse orientation. **i,** Distribution of average trajectories in the horizontal plane (n = 1 ganglion, stained with acTUBA). Color key corresponds to nerve target location in the horizontal plane. There is a bias away from the proximal-distal axis. Key illustrates the horizontal orientation. acTUBA, acetylated alpha tubulin; CB, cell bodies; ExS, external side; InS, internal side; NP cap, neuropil cap; SG, sucker ganglion.

We also examined how the trajectories of the nerve fibers vary according to root location (Fig. 8h, i). We found the distribution of trajectories follows the distribution of the roots (Fig. 8j). The nerves issuing from the aboral territory between the aboral and lateral gaps extend aborally in the direction of the aboral ACBM and the oral roots leading to the ANC (Fig. 8h, j). Those issuing from the oral territory extend laterally and orally in the direction of the lateral ACBM, sucker’s acetabulum, and ANC oral roots targeting the sensory epithelium of the sucker (Fig. 8h, j). These results establish a spatial regionalization to nerve fiber trajectories along the aboral-oral axis of the sucker ganglion.

In the ANC, there is an asymmetry in nerve fibers targeting the sucker: those issued from the external side of the ANC cover more territory of the sucker compared with those issued from the internal side (Olson et al., 2025). We asked if such an asymmetric in nerve fiber distribution is present in the sucker ganglion by examining the average trajectories along the external-internal axis (See Fig. 3a). We did not detect a similar kind of asymmetry in the sucker ganglion nerve fibers, with the external and internal sides similarly covered (Fig. 8j, k). We found instead a bias away from the proximal-distal axis and towards the external-internal axis, proposing a variation in function between the proximal-distal and external-internal planes (Fig. 8k). These results also demonstrate a divergence between sucker ganglion and ANC nerve fiber distributions, signifying that the sucker ganglion has processing roles distinct from those of the ANC.

## Discussion

The sucker ganglia are an exceptional feature of the octopus nervous system. First, across the eight arms of an adult *O. bimaculoides*, there are estimated to be over 1000 ganglia. Second, as noted by Graziadei (1971) and confirmed in the results of this study, they have an inverted structure. Third, as we document here, they have a rich internal complexity, an aboral inner belt with neuronal cell bodies, an oral outer belt devoid of neurons, and a connective tissue plate subdividing the ganglion as a whole (Fig. 9a).

**Figure 9:**
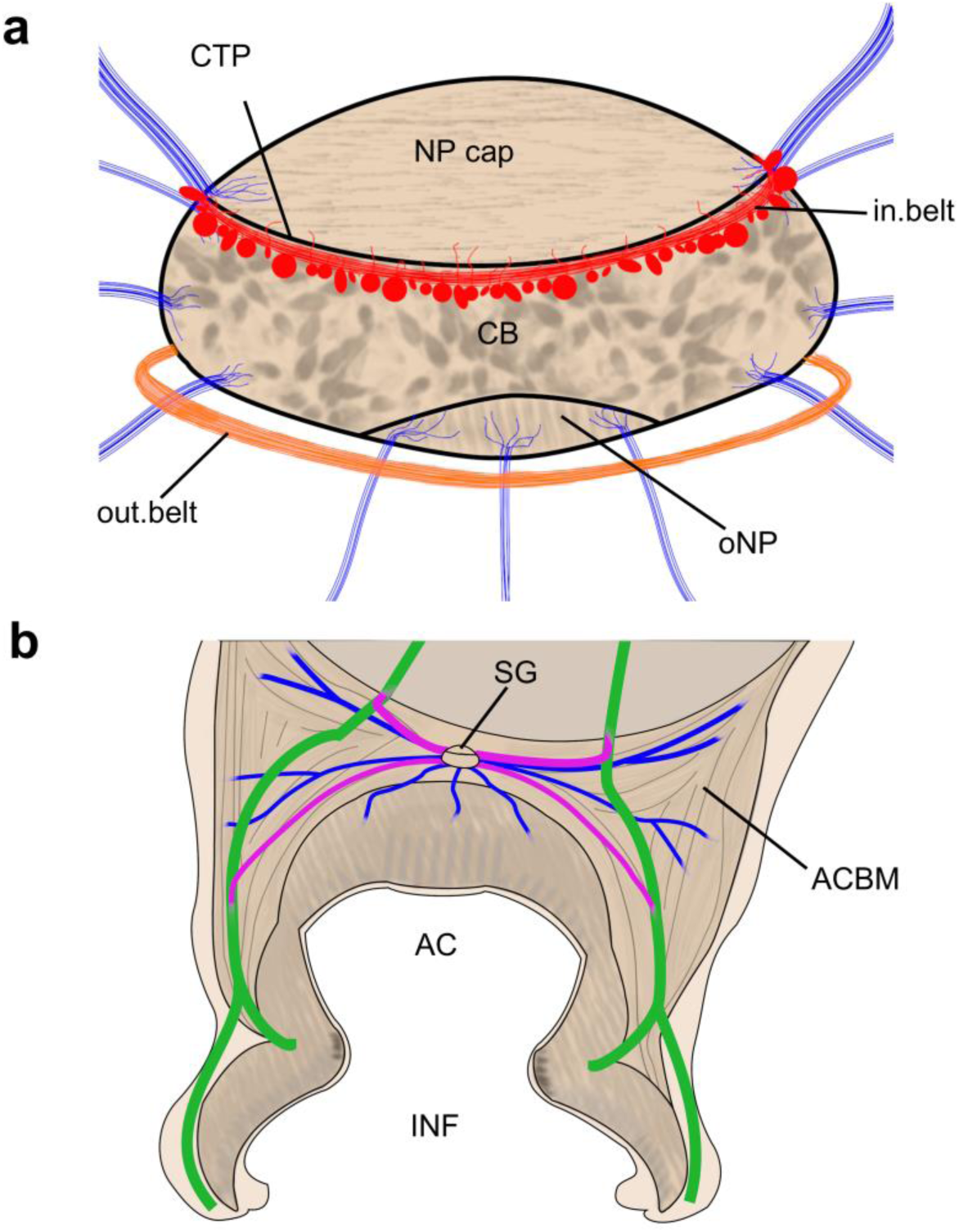
Summary of sucker ganglion internal complexity and nerve fiber connections. **a,** Cartoon depicting internal organization of the sucker ganglion (SG). The inner belt (red; in.belt), outer belt (orange; out.belt) and nerves (blue) connecting to the SG are highlighted. **b,** Schematic depicting nerve connections. Oral nerves from the axial nerve cord are green. Nerves from the SG that target the oral nerves are pink. Nerves from the SG that target local musculature are blue. AC, acetabulum; ACBM, acetabular brachial muscles; CB, cell bodies; CTP, connective tissue plate; in.belt, inner belt; INF, infundibulum; NP cap, neuropil cap; oNP, oral neuropil; out.belt, outer belt; SG, sucker ganglion.

Our molecular investigations have shown positive expression for both motor and sensory neuron markers, indicating a dual sensory and motor role for the sucker ganglion. The molecular profile of the sucker ganglion, however, is less complex than that of the ANC, consistent with a divergence in function between these two structures. We failed to detect positive expression of *DRGX* within the ganglion, although both the CBL of the ANC and the ACBM are enriched with *DRGX*-expressing cells. Vertebrate mammalian studies suggest that *DRGX* is expressed in nociceptive sensory cells (Chen et al., 2001; Rebelo et al., 2006). Primary nociceptive cell bodies for the sucker, therefore, may be situated within the ANC or the sucker musculature but not in the sucker ganglion. Our positive detection of *PIEZO* within the sucker ganglion and ACBM support Graziadei’s (1965) observation of muscle receptors. This molecular evidence suggests that the sucker ganglion carries proprioceptive information for the sucker.

The presence of both sensory and motor cells suggests that the sucker ganglion is well suited to modulate local reflexes for the sucker without input from the ANC. Indeed, the sucker ganglion extends nerve fibers to the local musculature and cells within the muscles feed back to the sucker ganglion. Classic literature has also suggested the presence of sensory-motor cells extending from the sucker ganglion: a single cell body with one sensory projection detecting muscle stretch and one motor projection to activate muscle (Martoja and May, 1955). Future work is needed to investigate this interesting possibility.

The data strongly imply functional differences in sensory-motor processing across the 3 cardinal axes for a sucker (see Fig. 3). The sucker ganglion sends nerve fibers in both the oral and aboral directions towards the oral roots issued from the ANC to the sucker epithelium. The connections orally suggest that the sucker ganglion may also exchange information with the sensory epithelium. The connections aborally, combined with the presence of large nerve bundles, suggest that the sucker ganglion sends information to and receives information from the ANC. Hence, the sucker ganglion is situated for both local reflexes and relays to the ANC (Fig. 9b).

There is also a bias of nerve fibers towards the external-internal axis and away from the proximal-distal axis. Along with asymmetric *FMRF* gene expression, this could underlie preferential sensorimotor activity along the external-internal axis. Further studies of the distribution of specific chemoreceptor subtypes around the sucker’s sensory epithelium could elucidate any biases in sensory transduction, and an examination of individual sucker movements along the cardinal axes could uncover biases in motor action. Interestingly, many behaviors utilizing multiple suckers, such as passing food along or grasping an object, require the movement of individual suckers along the proximal-distal axis (Wells, 1964; Altman, 1968; Sivitilli et al., 2022). We did not detect any direct connection between sucker ganglia, so the ANC most certainly handles long distance connections and coordination of motor action between suckers (Rowell, 1963; Graziadei, 1971; Olson et al., 2025). The ANC may also rely on its own separate circuits for control of coordinated motor actions between suckers or instead coordinate the activity of the sucker ganglia. Functional studies are needed to assess the relative roles of the ANC and the sucker ganglion for motor control of the suckers.

Structure and function of peripheral ganglia vary greatly across phyla (Tauc, 1967). Vertebrate sympathetic autonomic ganglia, for example, are traditionally thought of as relay stations for faithful transmission of central nervous system signals to effectors (Tauc, 1967; Nauta and Feirtag, 1986; Karemaker, 2017). Our data suggest that the sucker ganglia do not have an analogous function, but rather exhibit substantial internal integrative circuitry. A comparison to other peripheral ganglia in cephalopods is also informative. Cephalopods have a pair of stellate ganglia located in the mantle, most famously known for the giant fiber system in squid (Young, 1936, 1972; Hodgkin and Huxley, 1939; Lund, 1971). These ganglia are implicated in control of mantle contractions for respiration and escape as well as adaptive coloration through direct relays from the brain and local ganglionic processing (Young, 1938; Wilson, 1960; Lund, 1971; Bühler et al., 1975; Gonzalez-Bellido et al., 2014, 2018). Structurally, the stellate ganglia have a clear regionalization along the dorsal-ventral axis in the size of the cells and the structure of the NP (Lund, 1971; Young, 1972). This is reminiscent of the regionalization we report in the sucker ganglion along the aboral-oral axis. However, unlike the sucker ganglion, the stellate ganglia have a typical ganglionic structure of cell bodies wrapping neuropil in its center (Lund, 1971; Young, 1972). The stellate ganglia also do not have an equivalent outer or inner belt like the sucker ganglia. Thus, even in the context of other cephalopod ganglia, the structure of the sucker ganglion is strikingly unique.

## Conclusion

The sucker ganglion is embedded in the sucker musculature, strategically situated between the ANC and the acetabulum of the sucker. Previous anatomical studies have proposed, in turn, a sensory, a motor, or a combined sensory and motor role for the ganglion (Guérin, 1908; Martoja and May, 1955; Rowell, 1963; Graziadei, 1965; Graziadei and Gagne, 1976). Our molecular analysis does point to the sucker ganglion having populations of both sensory and motor neurons. Having intermixed sensory and motor neurons does not, however, guarantee an integrative role for the ganglion. It could simply serve a bidirectional relay structure for signals traveling to and from the ANC. It is our cellular analysis demonstrating a complex internal organization and a suite of fiber projections independent of those of the ANC to the sucker apparatus (Fig. 9) that lead us to conclude that the sucker ganglion has specialized and unique integrative roles in sucker sensory-motor neuronal processing.

## Acknowledgements

We thank Dr. Chuck Winkler of Aquatic Research Consultants for providing us with octopuses, and Dr. Caroline Albertin, Dr. Rahul Parnaik, and Ms. Natalie Grace Schulz for cDNAs. We thank Ms. Natalie Grace Schulz for the Picrosirius Red staining protocol and for comments. Imaging was performed at the University of Chicago Integrated Light Microscopy Core (RRID: SCR_019197). We extend special thanks to Dr. Christine Labno and Mr. Khalil Rodriguez for their invaluable assistance. We thank Ms. Caroline Miller for assistance with whole mount image processing. This work was supported by the NIH UF1NS115817 award (CWR).

---

**Supplemental Video 1: 3D-rendering of the sucker ganglion.** Video depicts 3D view of the sucker ganglion stained with acetylated alpha tubulin (acTUBA) and presented in Figure 3. Note that in 3D, the inner and outer belts are spatially segregated.

